# HECIL: A Hybrid Error Correction Algorithm for Long Reads with Iterative Learning

**DOI:** 10.1101/162917

**Authors:** Olivia Choudhury, Ankush Chakrabarty, Scott J. Emrich

## Abstract

Second-generation sequencing techniques generate short reads that can result in fragmented genome assemblies. Third-generation sequencing platforms mitigate this limitation by producing longer reads that span across complex and repetitive regions. Currently, the usefulness of such long reads is limited, however, because of high sequencing error rates. To exploit the full potential of these longer reads, it is imperative to correct the underlying errors. We propose HECIL—Hybrid Error Correction with Iterative Learning—a hybrid error correction framework that determines a correction policy for erroneous long reads, based on optimal combinations of decision weights obtained from short read alignments. We demonstrate that HECIL outperforms state-of-the-art error correction algorithms for an overwhelming majority of evaluation metrics on diverse real data sets including *E. coli*, *S. cerevisiae*, and the malaria vector mosquito *A. funestus*. We further improve the performance of HECIL by introducing an iterative learning paradigm that improves the correction policy at each iteration by incorporating knowledge gathered from previous iterations via confidence metrics assigned to prior corrections.

**Availability and Implementation:** https://github.com/NDBL/HECIL

## 1 Introduction

Current advances in next-generation sequencing (NGS) have fueled genomics-driven research by inexpensively generating highly accurate ‘reads’ or DNA sequence fragments. Second-generation sequencing technologies, for example Illumina [1] and 454 pyro-sequencing [2], generate short reads that are sometimes not ideal for downstream applications such as assembling complex genomes [3]. To ameliorate this issue, thirdgeneration sequencing techniques introduced by Pacific Biosciences [4, 5] and Oxford Nanopore [6, 7, 8] generate significantly longer reads. These long reads typically contain thousands of base-pairs (see for example [9]) and are not subject to amplification or compositional biases often exhibited by second-generation sequencing [10]. Long reads also overcome issues associated with repetitive regions and large transcript isoforms. In spite of these significant advantages, a critical limitation of long reads produced by thirdgeneration sequencing methods is that they generally exhibit high error rates: for example, up to 20% error has been reported using PacBio [11, 12], and 35% using Oxford Nanopore [13].

Various correction algorithms have been proposed for reducing the currently high error rates prevalent in long reads. For example, HGAP [14] is a self-correcting algorithm (that is, it does not rely on additional sequencing data) that performs correction by computing multiple alignments of high coverage long reads. Another class of correction algorithms rely on short reads generated from the same (or related) samples, and is therefore referred as *hybrid correction algorithms*. For instance, the authors in [13] proposed the Nanocorr algorithm in which high-quality Illumina MiSeq reads are used to correct Oxford Nanopore reads.

Popular hybrid correction algorithms for PacBio data include: LSC [3], PacBioToCA [9], LoRDEC [15], proovread [16], and CoLoRMap [17]. Most of the methods listed here do not systematically utilize localized information such as base quality of the short reads or variant information between individuals. The importance of incorporating base quality in correcting noisy sequencing data, for example, has been emphasized in [18].

This motivates our proposed algorithm, HECIL: a novel hybrid error correction framework that computes a correction policy by selecting an optimal combination of decision weights based on base quality and mapping identity of aligned short reads. We compare HECIL’s performance to existing hybrid correction algorithms on real prokaryotic and eukaryotic data and, for an overwhelming majority of the evaluation metrics, show that HECIL outperforms its competitors. We also propose an iterative learning framework that improves HECIL’s correction accuracy both in terms of alignment and assembly-based metrics by incorporating knowledge derived from high-confidence corrections made in prior iterations. We speculate that the proposed iterative learning formalism can be incorporated into other contemporary hybrid error correction algorithms to improve performance, provided that each iteration does not require prohibitive execution time.

## 2 Methods

HECIL works under the standard assumption that all reads are derived from highly similar individuals. We begin by aligning the given set of short reads to the long reads. For each alignment, we compute normalized weights using base quality information and alignment identity of the underlying short reads. The short read that maximizes the sum of these normalized weights is used for correction. In this manner, we tend to select higher quality short reads that have a suitable degree of overlap with a long read. This forms the *core algorithm* of HECIL.

Next, we optionally define a subset of these corrections as “high confidence” and correct only these high-confidence errors. By introducing elitism to the correction procedure based on confidence, the updated long reads now exhibit slightly higher consensus (or similarity) with the short reads. Therefore, we expect to obtain slightly higher quality alignments for fixing lower confidence corrections in subsequent iterations: this is the intuition behind the iterative learning procedure. Herein, we discuss each of these steps in detail.

### 2.1 HECIL’s Core Algorithm

#### 2.1.1 Step 1: Quick Correction

As recommended in [16] and [17], we first obtain read alignments using BWA-MEM [19] and then mark positions with disagreements (for example: mismatches, insertions, and deletions) on long reads as *questionable*. For each questionable position on the long read, we investigate the set of short reads that align to it. If there is strong consensus (determined by a threshold 0 *" η ≤* 1 selected by the user), we replace the questionable base on the long read with the respective aligned base of the short read. This *quick correction* step is illustrated in Fig. 1-(A). This step is inspired by the majority voting method proposed in [3]. However, contrary to corrections based on a simple majority, we adopt a stricter threshold of at least 90% consensus (*η* = 0.9) to be eligible for quick correction. Shifting from majority voting to strong consensus prevents spurious corrections made on the basis of high-frequency, low-quality short reads. Note that quick correction also reduces the search space in the next step of HECIL’s core algorithm.

**Figure 1:**
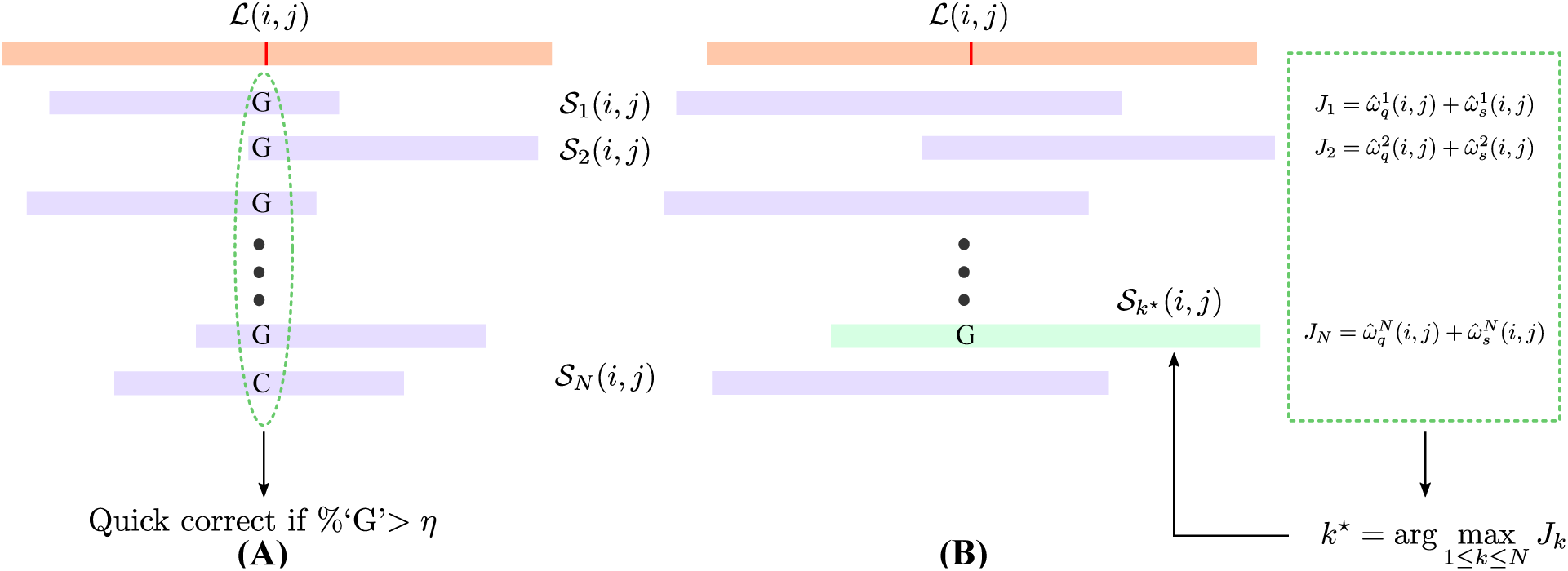
Illustration of HECIL’s core algorithm. The orange rectangle denotes an erroneous long read and the purple rectangles represent aligned short reads. (**A**) Quick correction with high consensus. (**B**) Optimization-based correction: The green dashed box depicts the objective function values, from which the optimal short read (green rectangle) is selected for correction.

#### 2.1.2 Step 2: Optimization-based Correction

For the remaining questionable bases, we employ an optimization-based correction framework. Let *ℒ*(*i, j*) be the *j*th questionable base corresponding to the *i*th long read. Suppose *N* short reads align to this *ℒ* (*i, j*);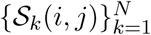denotes the set of aligning short reads. For each *k* = 1, 2*,…, N* we assign two normalized weights 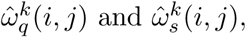 and 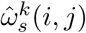,representing the quality and similarity of the *k*th short read, respectively. The normalized quality weight is given by

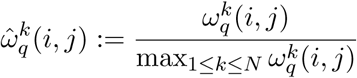

where the scalar *ω*_*q*_(*i, j*) is determined by extracting the PHRED quality score readily available from FASTQ files. The normalized similarity weight 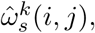 is obtained by calculating the alignment identity, defined as the number of exact matches of the *k*th short read *S*_*k*_(*i, j*) to the long read *ℒ*(*i, j*), divided by the length of *S*_*k*_(*i, j*). Untrimmed short reads, therefore, may result in a lower estimated 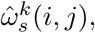, which is why we adhere to trimmed short reads in this study. For each short read, we compute a cost by taking a convex combination of the two normalized weights

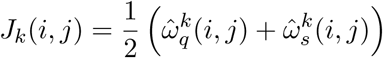

We then solve the following optimization problem:

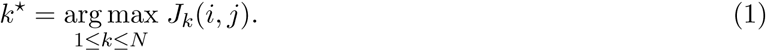

which yields the index *k*^*\**^ of the short read *S*_*k*_*\** (*i, j*) that exhibits the maximum combined quality and similarity weight. In case there is a conflict amongst maximizers, the short read with highest quality is selected to be the winner. Note that the optimal cost for each *ℒ* (*i, j*) is denoted by *J*_*k*_*\** (*i, j*). Subsequently, we replace the erroneous base *ℒ* (*i, j*) on the long read with the corresponding base of the short read *S*_*k*_*\** (*i, j*). This procedure is illustrated in Fig. 1(B).

If perfect consensus (that is, *η* = 1 in Step 1) is reached amongst all the short reads, there is no need to perform Step 2, because both steps will yield identical corrections. Similarly, if we select a consensus threshold *η ∈* (0, 1), then the probability that the quick correction value matches the optimization-based correction value is *η*, irrespective of the cost function selected. Therefore, choosing *η* close to 1 ensures that quick correction matches optimization-based correction with high-probability. We do not set *η* strictly equal to 1 hypothesizing that achievement of perfect consensus is rare in practice.

Also note that the quality of a short read and its alignment identity with the long read are not contending objectives. That is, a high quality read does not always imply low similarity and vice versa. Therefore, we consider a convex combination of these objectives as in equation (1) rather than formulating a multi-objective optimization problem and searching for Pareto-optimal solutions.

### 2.2 Improving Correction Performance via Iterative Learning

A definition of iterative learning that closely resembles our proposed approach in this paper is found in [20]: iterative learning “considers systems that repetitively perform the same task with a view to sequentially improve accuracy”. Here, the *same task* refers to the core algorithm of HECIL, and the goal is to improve error corrections in the *l* th iteration by learning from high-confidence corrections in the (*l -* 1)th iteration (see Fig. 2). Unlike the iterative proovread algorithm, our proposed iterative scheme leverages confidence metrics assigned to prior corrections for informing later decisions.

**Figure 2:**
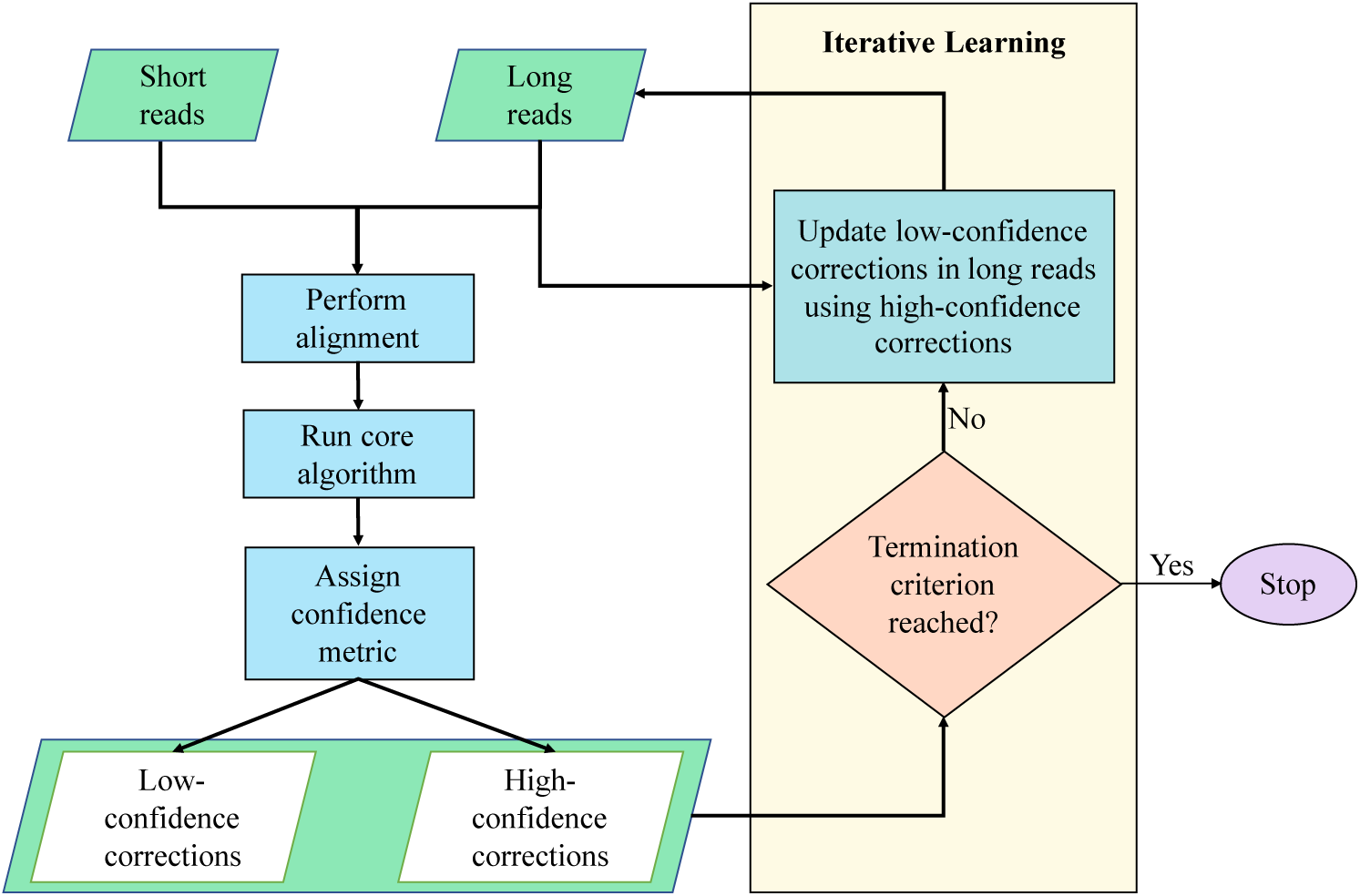
Iterative learning procedure with the HECIL core algorithm for error correction. Alternative hybrid-error correction algorithms could seamlessly replace the core algorithm.

#### 2.2.1 Assignment of confidence

For each ℒ (*i, j*) in the *l* th iteration, suppose the corresponding optimal cost obtained by solving (1) be denoted by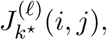, and let *μ*^(*l*)^ denote the α -percentile computed over all these optimal costs. Here we select α *>* 0.85 so that a small percentage of the optimal corrections are considered to be of high confidence. Selecting a high value of α ensures that only the highest quality corrections will always inform future iterations. Adversely, selecting *α* too close to one will result in slower improvement of correction accuracy, because large α implies that very few corrections are deemed high confidence. Therefore, the increment in information used to update the correction policy in the following iteration will be limited.

#### 2.2.2 Realignment based on high-confidence corrections

We *learn* in successive iterations by re-aligning the updated long reads to the short reads. Note that, for each iteration, the updated context of ℒ(*i, j*) could generate entirely different sets of aligned short reads, as well as disparate localized information from previous iterations, leading to the calculation of different sets of normalized weights 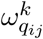 ans 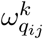. This why the confidence threshold *μ*^(*l*)^ is recomputed based on the statistics of the optimal costs (namely, the percentile measure) and not fixed. The sites on the long read corresponding to low-confidence short reads are left to be changed via the core algorithm in a subsequent iteration while the high confidence changes in prior iterations are effectively fixed.

#### 2.2.3 Termination criteria

We propose two criteria for termination. In the first criterion, if the fractional reduction in the number of unique *k* -mers (overlapping substrings of length *k*) between two successive iterations is below a given threshold *ε ∈* (0, 1). That is,

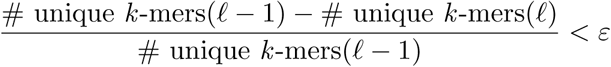

then we terminate after the *l* th iteration. Specific arguments why *k* -mers are used for termination are provided in Section 3.2.1.

The second criterion involves pre-calculating the number of iterations to achieve at least *M* ^*′*^ highconfidence corrections. At each iteration, (1 *-α*) *×* 100% of the corrections are retained (by definition of the *α*-percentile), implying that *αM* errors remain for the next iteration. Thus, the number of lowconfidence corrections remaining after *n* iterations will be *α*^*n*^*M*. If the user requires at least *M* ^*′*^ highconfidence corrections to be made iteratively, one needs to select *n* that satisfies *M* ^*′*^ *<* (1 *-*α^*n*^)*M*. In such a case, algebraic manipulations and taking logarithms yield the following bound on the required number of iterations

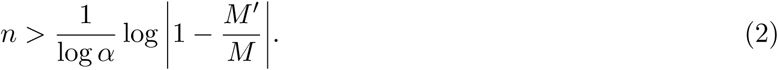

Note that both logarithm terms are negative. Based on the above, after *n* corrections, where *n* satisfies (2), we expect that at least *M* ^*′*^ high confidence corrections will be made.

## 3 Results

All experiments in this section were run on Dell PowerEdge R815 servers with AMD Opteron processor 6378, Quad 16 core 2.4 GHz CPU, 32 cores, 512 GB RAM, and 2 x 300 GB 15K RPM SAS drives. We use the Unix time command to record the runtime and memory usage of each tool.

### 3.1 Data acquisition

We test the performance of HECIL on three datasets of varying size: the bacterial genome of *E. coli*, the fungal genome of *S. cerevisiae*, and the malaria vector genome of *A. funestus*. We explore PacBio-sequenced long reads, Illumina-sequenced short reads, and reference genomes of *E. coli* and *S. cerevisiae*, as suggested in the supplementary material of [17]. Long reads of *E. coli* are filtered to exclude reads shorter than 100 bp. The final set contains 33,360 reads that total 98 million bases (Mbp). The corresponding short reads (accession ID: ERR022075) comprise 22,720,100 reads that are 202 bp long. The strain K-12 substr. MG1655 is used for our alignment-based validation of HECIL. To test *S. cerevisiae* data we use 1,758,169 long reads, with 1,402 Mbp and 4,503,422 short reads (accession ID: SRR567755). The reference genome of strain S288C is 12.2 Mbp in size. We obtain long reads for *A. funestus*^1^, comprising data from 44 flowcells, ranging between 59,937 and 244,754 reads. Due to the high computational time required by proovread and CoLoRMap to correct the reads of all flowcells, we present a comparative analysis based on a random selection of three: flowcells 1, 4, and 16. Short sequences (accession ID: SRR630594) consists of 37,797,235 reads. The reference genome of strain Fumoz (GenBank assembly accession: GCA 000349085.1) is used for validating corrections.

### 3.2 Evaluation metrics

#### 3.2.1 *k* -mer-based

We employ the widely-used *k* -mer counting tool Jellyfish [21] to compute the number of unique *k* -mers obtained after each correction algorithm is executed. Since errors in long reads are uniformly distributed across their length, large numbers of uncorrected errors often greatly inflate the number of unique *k* -mers observed. Further, the authors in [22] reported that the set of common *k* -mers between the highly accurate short reads and the erroneous long reads were crucial in improving the quality of data for downstream analysis. Therefore, a correction algorithm that reduces the number of unique *k* -mers while increasing the number of valid *k* -mers is desirable. Fig, 3 gives an illustrative example of this idea based on one *A. funestus* flowcell.

#### 3.2.2 Alignment-based

After each method of correction, we align corrected long reads to its reference genome using BLASR [23]. In addition to computing the number of aligned reads and aligned bases, we evaluate matched bases, that is, the ratio of total number of matched bases and length of sequences in the long reads. We calculate percent identity (PI) by the ratio of matches to alignment length. Since PI is usually above 90%, we also determine the number of aligned reads with PI above 90%.

#### 3.2.3 Assembly-based

One of the most important downstream applications of long reads is *de novo* genome assembly. For this purpose, we use the assembler Canu [24], specifically designed for noisy long reads. We then use QUAST [25] to evaluate the quality of the resulting assembly. We measure total number of contigs, length of the longest contig, and total length, i.e., total number of bases in the assembly. We report the values of N50 (minimum length such that contigs of that length or longer consists half the assembly), and NG50 (minimum length such that contigs of that length or longer consists half the reference assembly). As suggested in [24], we further measure accuracy by aligning the assembled genome to the reference genome using MUMMer’s dnadiff tool [26]. In this context, we compute percent of aligned bases (with respect to reference and query) and average identity of 1-to-1 alignment blocks (with respect to reference and query).

**Figure 3:**
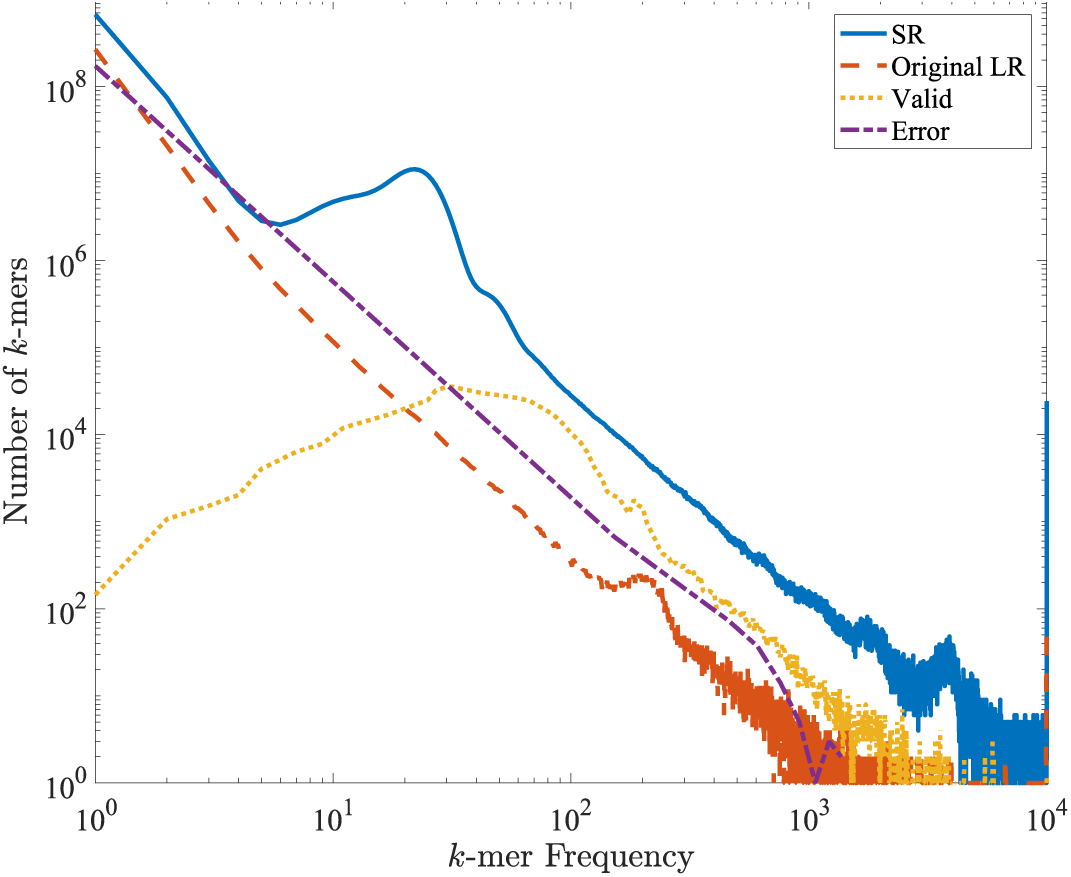
Distribution of *k* -mer frequency (*k* =17) in *Anopheles funestus* flowcell #16. The *x* and *y*-axes denote *k* -mer frequency and count of frequency, respectively. The blue line and dashed red line represent *k* -mers generated from short reads (SR) and original long reads (Original LR), respectively. As discussed in Section 3.2.1, the dotted yellow line indicates that majority of the valid *k* -mers have high frequency. The purple dot-dashes, representing error *k* -mers (not found in short reads), mostly consists of unique *k* -mers.

### 3.3 Comparative analysis

We compare the performance of HECIL with cutting-edge hybrid error correction tools such as proovread2.14.0, LoRDEC-0.6, and CoLoRMap. We use the above-mentioned *k* -mer-based, alignment-based, and assembly-based metrics to assess the performance of each approach. The comparative results for *k* -merbased and alignment-based parameters are presented in Table 1. We report the parameters before correction (original) and after each method of error correction.

**Table 1:**
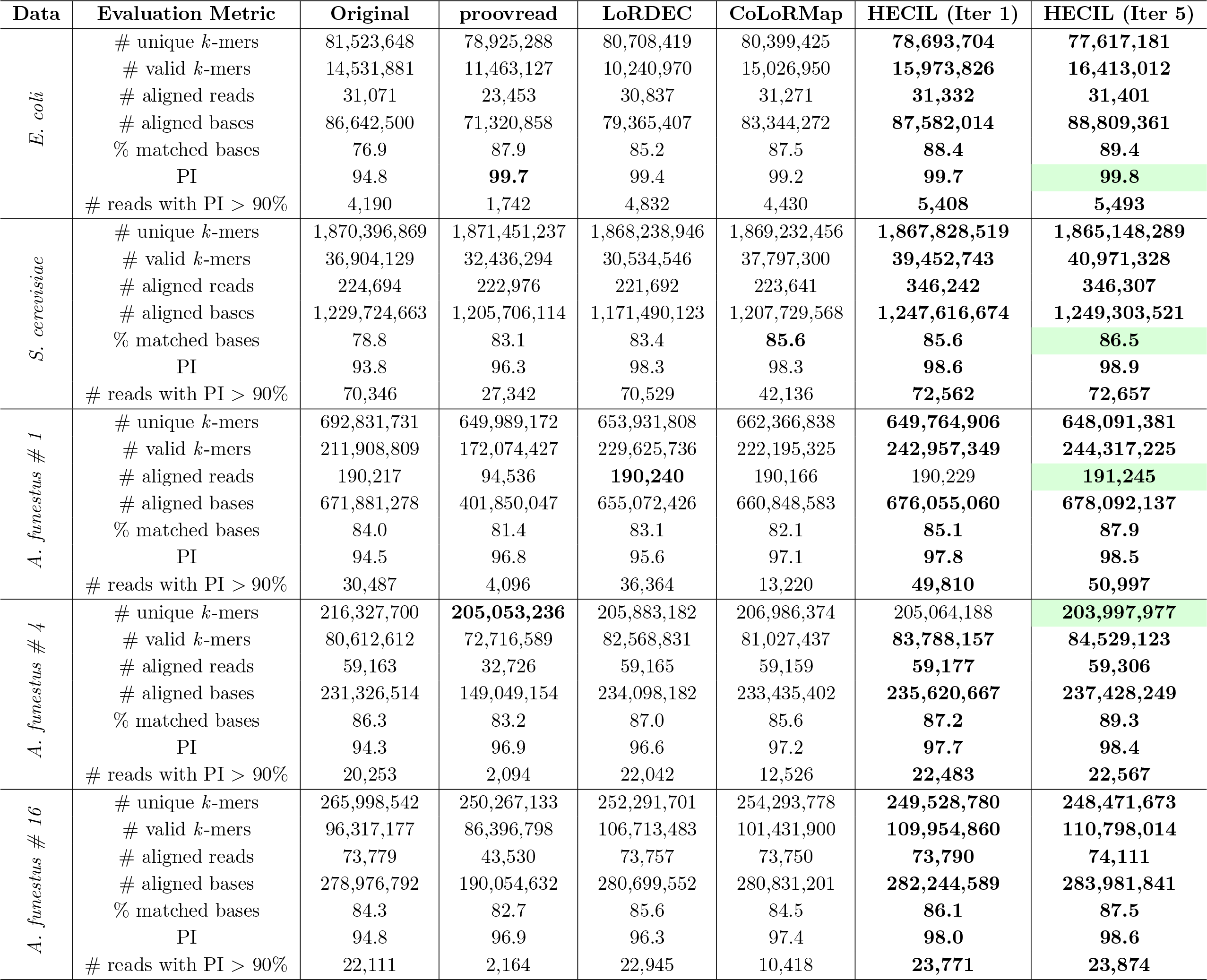
Comparison of alignment-based metrics obtained from testing *E. coli*, *S. cerevisiae*, and *A. funestus* on proovread, LoRDEC, CoLoRMap, and HECIL. For the case of HECIL, metrics are reported before and after using the iterative learning algorithm; specifically, iteration 1 (the core algorithm) and iteration 5 (with four rounds of learning) are shown. The colored cells in the table denote metrics where the core algorithm did not outperform its contemporaries, but with iterative learning, achieved the best performance.

As discussed in [17], CoLoRMap performs better than proovread and LoRDEC, when tested on *E. coli* and *S. cerevisiae*. Long reads corrected by the core algorithm of HECIL (iteration 1) generate the lowest number of *k* -mers, with the exception of the data set *A. funestus* flowcell 4, where it is still comparable to the best results obtained from proovread. A corresponding increase in valid *k* -mers indicates higher consensus to the accurate short reads, hence higher accuracy of the corrected long reads. For all data sets, HECIL consistently produces more valid *k* -mers. It also results in highest number of aligned bases and reads, highest value of percent identity, and reads with at least 90% percent identity to its reference.

We present the results of assembly-based metrics in Table 4. For *E. coli* and yeast, HECIL generates more contiguous assembled long reads, compared to the existing tools, except CoLoRMap where there are ties. The size of the longest contig and the number of bases in the assembled data are highest for our proposed approach. The standard assembly quality parameters like N50 and NG50 have highest values after using HECIL for correction. Furthermore, the assembled genomes of HECIL have higher aligned bases and 1-to-1 alignment identity. Since we use a subset of mosquito’s flowcell data, the proportion of aligned bases in the reference genome with respect to the query genome is low.

For highly heterozygous samples such as insects like mosquitoes [27], low frequency bases in aligned short reads may indicate inherent variation that are not necessarily sequencing errors. Correction algorithms that solely rely on a consensus call or majority vote often discard these heterogenous alleles. The optimization in Step 2 of HECIL is not biased by bases which have high frequency, and hence, is better able to capture variation between similar individuals. This is corroborated by the results obtained from testing HECIL on the highly heterozygous mosquito data set of *Anopheles funestus*.

Although the performance of hybrid correction algorithms largely depend on the set of high coverage short reads, we devise additional experiments to verify that restraining the coverage of short reads does not have a deleterious effect on HECIL. We down-sample short reads by randomly selecting 50%, 25%, and 12% of the data to be used for correction. In *E. coli*, this results in a subset of short reads for correction with an average coverage of 62*×*, 33*×*, and 18*×*, respectively. In Table 2, we present *k* -mer-based and alignment-based parameters from correcting long reads of *E. coli* with the down-sampled short reads using HECILand assembly-based parameters from the lowest coverage (18*×*; Table 4). Thus, HECIL shows potential for use in projects that do not have high coverage short read data readily available: this is especially important in larger eukaryotic genomes sequenced predominantly with longer read technology.

**Table 2:** Comparison of alignment-based metrics with downsampled *E. coli* short reads using HECIL’s core algorithm.

| Evaluation Metric | All SRs | 50% SRs | 25% SRs | 12% SRs |
| --- | --- | --- | --- | --- |
| # $k$-mers | 84,294,561 | 84,121,231 | 83,986,262 | 83,909,706 |
| # unique $k$-mers | 78,693,704 | 78,292,463 | 78,097,941 | 78,008,319 |
| # valid $k$-mers | 15,973,826 | 15,889,155 | 15,737,641 | 15,576,317 |
| # aligned reads | 31,332 | 31,328 | 31,322 | 31,318 |
| # aligned bases | 87,582,014 | 87,359,227 | 87,288,475 | 87,196,236 |
| % matched bases | 88.4 | 88.4 | 88.3 | 88.3 |
| PI | 99.7 | 99.7 | 99.7 | 99.6 |
| # reads with PI > 90% | 5,408 | 5,358 | 5,298 | 5,223 |

In Table 3, we compare the runtimes and maximum memory usage incurred in correcting each data set (see Methods). proovread, LoRDEC, and CoLoRMap were run with 16 threads. The workload of HECIL was split into 16 concurrent tasks, which were run in parallel. Computation time of hybrid error correction methods is mainly dominated by the underlying steps of generating intermediate data, such as mapping short reads to the long reads. Similarly, LoRDEC and CoLoRMap construct a graph data structure, which demands high computational resources. LoRDEC, however, uses the efficient GATB library [28], which lowers the overhead (see Table 3). Although our tool incurs higher computation time than LoRDEC, it is consistently faster (generally almost twice as fast) than the other correction methods and generates overall higher quality corrected long reads without a significant increase in memory consumption.

**Table 3:** Comparison of runtime (hh:mm:ss) and maximum memory footprint (GB) for correcting long reads. Only the best and worst *A. funestus* single iterations are shown.

| Data | Method | Runtime | Memory |
| --- | --- | --- | --- |
| *E. coli* | proovread | 6:15:37 | 11.4 |
|  | LoRDEC | 38:53 | 6.2 |
|  | CoLoRMap | 2:48:23 | 28.9 |
|  | HECIL | 1:16:55 | 9.1 |
| *S. cerevisiae* | proovread | 20:54:15 | 14.5 |
|  | LoRDEC | 3:43:12 | 6.1 |
|  | CoLoRMap | 7:57:49 | 38.2 |
|  | HECIL | 5:14:09 | 11.2 |
| *A. funestus*<br>(Flowcell # 1) | proovread | 76:13:47 | 8.8 |
|  | LoRDEC | 35:08:13 | 3.1 |
|  | CoLoRMap | 90:50:12 | 23.4 |
|  | HECIL | 46:06:47 | 8.3 |
| *A. funestus*<br>(Flowcell # 4) | proovread | 36:32:25 | 7.3 |
|  | LoRDEC | 11:25:05 | 6.7 |
|  | CoLoRMap | 32:18:30 | 20.7 |
|  | HECIL | 17:38:01 | 6.9 |

**Table 4:**
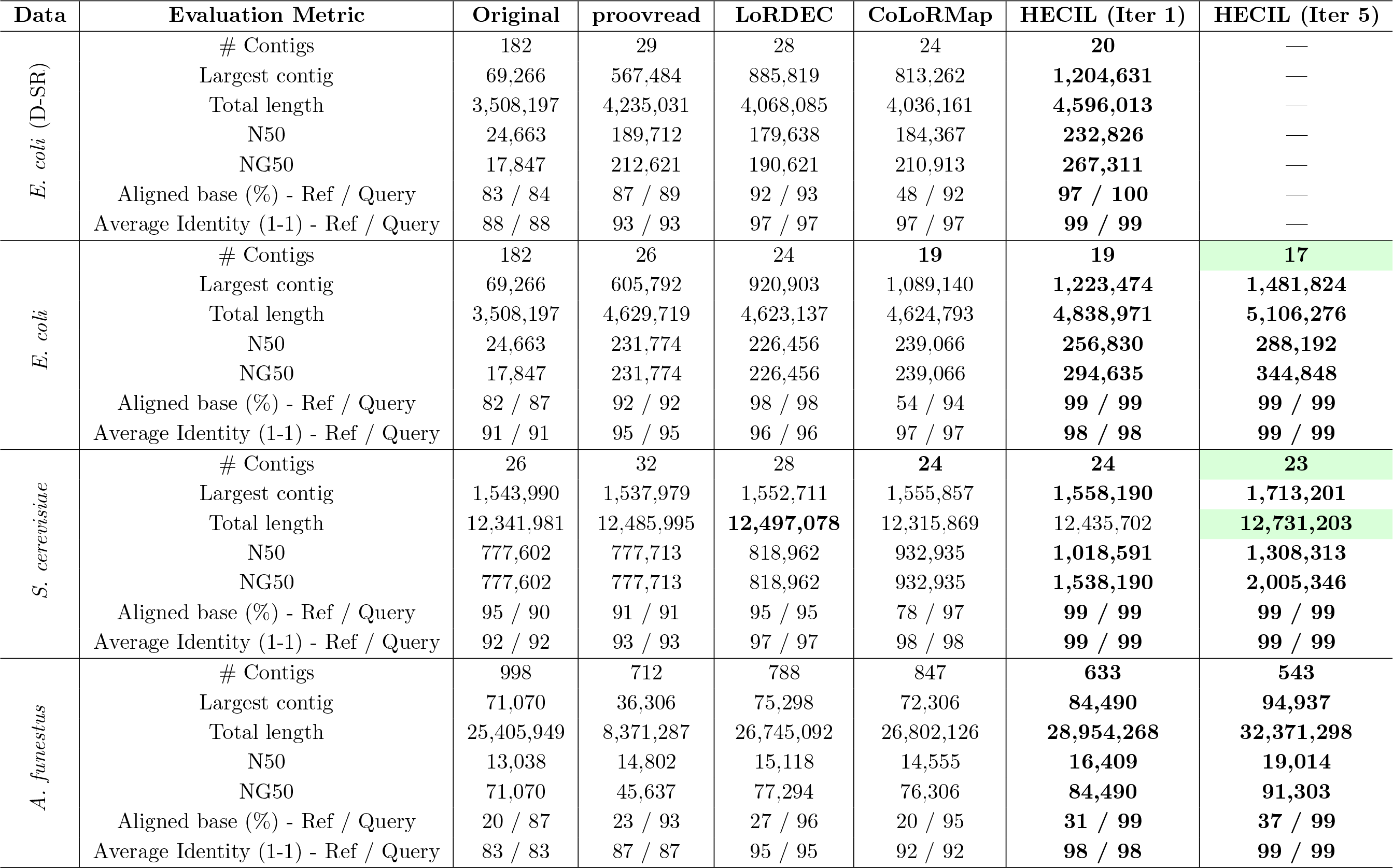
Comparison of assembly-based metrics obtained from testing *E. coli*: with downsampled short reads (D-SR) having 18x coverage (lowest coverage) and original short reads, *S. cerevisiae*, *A. funestus* (merged flowcells) on proovread, LoRDEC, CoLoRMap, and HECIL. For the case of HECIL, metrics are reported before and after using the iterative learning algorithm; specifically, iteration 1 (the core algorithm) and iteration 5 (with four rounds of learning) are shown. The colored cells in the table denote metrics where the core algorithm did not outperform its contemporaries, but with iterative learning, achieved best results.

### 3.4 Effect of Iterative Learning

We leverage our proposed iterative learning scheme on HECIL to demonstrate its effectiveness in further improving correction accuracy. We select a high-confidence cut-off of 85 percentile, that is, *α* = 0.85. The alignment-based incremental improvements obtained after each iterative correction of HECIL is presented in Fig. 4. For each data set (each column), we observe that the incremental metrics: number of fewer *k* - mers, number of additional aligned long reads, number of additional aligned bases, and additional percent of matched bases, improve after each iteration, until one of the termination criteria is reached. For the termination criteria, we select *ε* as 0.02 for the metric of unique *k* -mers and choose *M* ^*′*^*/M* = 0.5, which, using (2) results in at most *n* = 5 iterations. Based on this, we report alignment-based and assembly-based metrics obtained after the fifth iteration of HECIL in Tables 1 and 4, respectively. HECIL in conjunction with iterative learning consistently outperforms all the evaluation metrics. The colored cells in these tables indicate the parameters where the core algorithm of HECIL is comparable but does not outperform the alternatives, and the iterative version of HECIL consistently results in better performance. These results verify the utility of iterative learning-based extension of HECIL, particularly in heterozygous samples like the mosquito data set used in this study.

**Figure 4:**
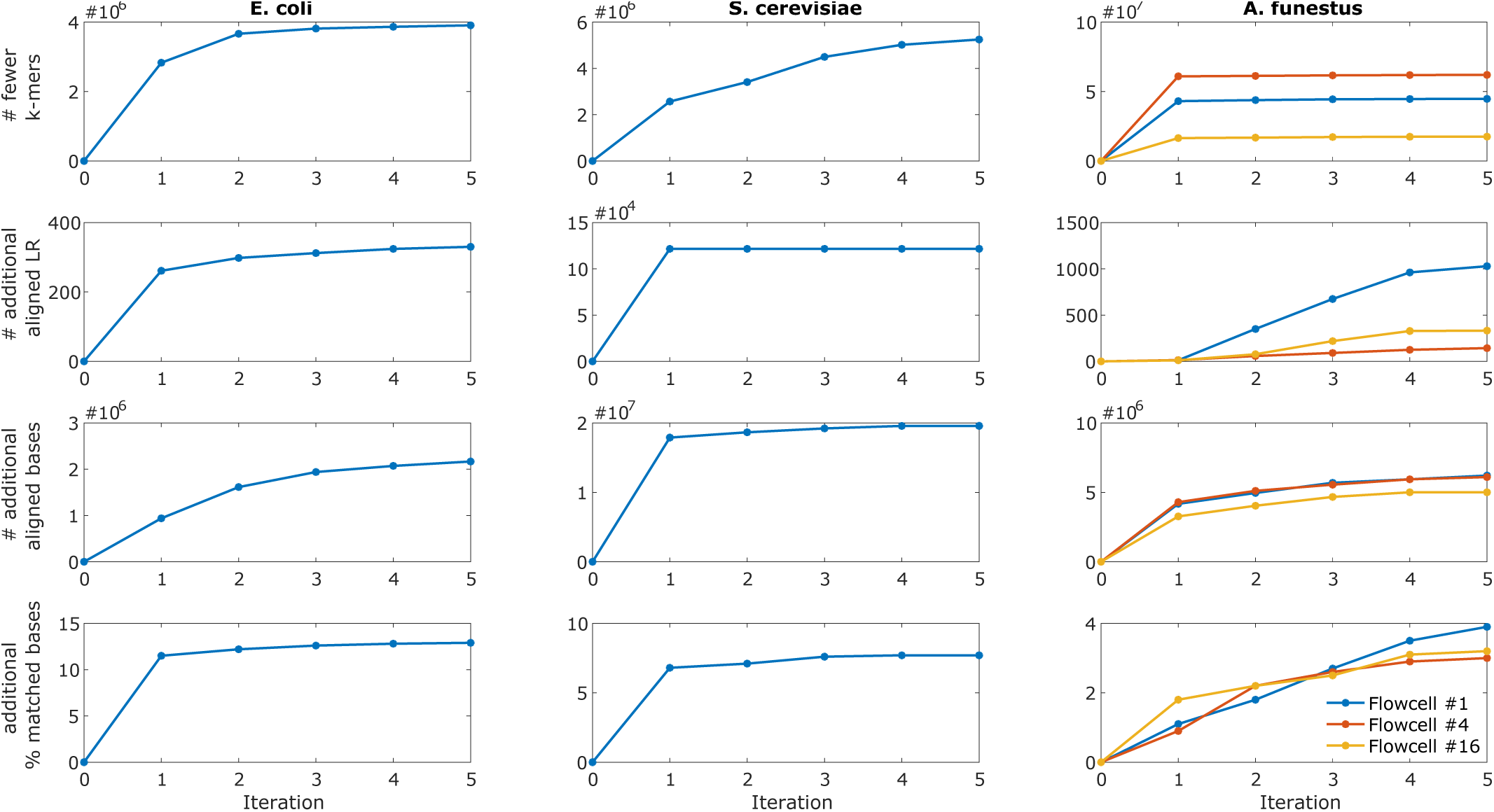
Improvement of alignment-based metrics (# fewer unique *k*-mers, additional aligned long reads, additional aligned bases, additional percent matched bases) for *E. coli*, *S. cerevisiae*, and *A. funestus* with iterative learning. The 0th iteration denotes the original data set and the 1st iteration indicates corrected data set obtained from running HECIL’s core algorithm.

## 4 Discussion

Third-generation sequencing techniques, particularly Single-Molecule Real-Time (SMRT) sequencing, is revolutionizing modern genomics. The usefulness of current long read data, however, is restricted due to high sequencing error rates. Hence, it is crucial to correct long reads prior to downstream applications like *de novo* genome assembly.

In this paper, we develop a novel approach of hybrid error correction called HECIL, which corrects erroneous long reads based on optimal combinations of base quality and mapping identity of aligned short reads. As seen in Tables 1 and 4, HECIL performs significantly better for an overwhelming majority of evaluation metrics, even with limited amounts of short reads available for correction.

To the best of our knowledge, this is the first time an iterative strategy for improving correction quality via informed realignment has been proposed. The method shows potential using the HECIL core algorithm, but could also be seamlessly integrated with other error correction algorithms that leverage short read alignments.

The current version of HECIL allows decomposition of the workload into independent data-parallel tasks that can be executed simultaneously. A natural extension of the tool is implementing multi-threading to achieve speedup on traditional machines.

## Acknowledgements

This work was supported by the Eck Institute for Global Health (EIGH) Ph.D. fellowship to Choudhury, and NIH R21AI123967 to Emrich.

## Footnotes

1 in process of submission to NCBI

